# Circuit robustness to temperature perturbation is altered by neuromodulators

**DOI:** 10.1101/178764

**Authors:** Sara A. Haddad, Eve Marder

## Abstract

In the ocean, the crab, *Cancer borealis*, is subject to daily and seasonal temperature changes. Previous work, done in the presence of descending modulatory inputs, had shown that the pyloric rhythm of the crab increases in frequency as temperature increases, but maintains its characteristic phase relationships until it “crashes” at extreme high temperatures. To study the interaction between neuromodulators and temperature perturbations, we studied the effects of temperature on preparations from which the descending modulatory inputs were removed. Under these conditions the pyloric rhythm was destabilized. We then studied the effects of temperature on preparations in the presence of oxotremorine, proctolin, and serotonin. Oxotremorine and proctolin enhanced the robustness of the pyloric rhythm, while serotonin made the rhythm less robust. These experiments reveal considerable animal-to-animal diversity in their crash stability, consistent with the interpretation that cryptic differences in many cell and network parameters are revealed by extreme perturbations.

## INTRODUCTION

Most animals must function in a variety of environments. Consequently, most neuronal circuits must be sufficiently flexible to allow for behavioral adaptation in response to environmental fluctuations. In the case of poikilothermic animals, environmental temperature fluctuations can be a strong evolutionary pressure that gives rise to robust mechanisms of temperature preference (Beverly et al., 2011; Hamada et al., 2008) that cause animals to physically translocate in their habitats to maintain relatively constant body temperature. Nonetheless, many animals, such as the North Atlantic crab, *Cancer borealis*, used in this study, experience substantial temperature fluctuations daily and seasonally (Donahue et al., 2009; Haeffner, 1977; Krediet and Donahue, 2009; Stehlik et al., 1991), even as they also move between deeper and shallower waters in search of their preferred temperatures.

Because temperature influences all biological processes to a different extent, it presents an extraordinary challenge to maintaining relatively stereotyped performance (Caplan et al., 2014; O’Leary and Marder, 2016; Rinberg et al., 2013; Robertson and Money, 2012; Roemschied et al., 2014; Schleimer and Schreiber, 2016; Tang et al., 2010; Tang et al., 2012). This is easily understood if one thinks of multiple membrane currents that contribute to some physiological process, each with a different temperature dependence (Partridge and Connor, 1978). Consequently, even modest temperature changes could cause significant changes or disruptions to that physiological system (Caplan et al., 2014; O’Leary and Marder, 2016). Nonetheless, there are examples of neurons and circuits that are well temperature compensated, although their individual membrane currents show appreciable temperature dependencies (Partridge and Connor, 1978; Roemschied et al., 2014).

Almost perfect temperature compensation is likely the exception, not the rule, and many poikilothermic animals show significant alterations in behavior as ambient temperature fluctuates (Soofi et al., 2014; Tang et al., 2010; Tang et al., 2012). Moreover, because all individuals of the same specie vary in the synaptic strengths and conductance densities found in individual neurons and networks (Goaillard et al., 2009; Golowasch et al., 1999; Lane et al., 2016; Norris et al., 2011; Roffman et al., 2012; Schulz et al., 2006; Schulz et al., 2007; Temporal et al., 2014; Wenning et al., 2014), it is not surprising that the population may show substantial variance in their sensitivity to temperature. In this paper, we use temperature as a perturbation of the activity of the crustacean stomatogastric ganglion (STG) to ask whether neuromodulation can stabilize network output in response to changes in temperature. To do so, we characterize state changes in network performance in response to temperature changes under a variety of modulatory conditions. The significant animal-to-animal variability revealed in the absence of strong modulatory drive impelled us to design new methods of analyzing this variability.

## RESULTS

The stomatogastric nervous system (STNS) has led to numerous insights into the basic mechanisms of the generation of rhythmic movements and their modulation (Marder, 2012; Marder and Bucher, 2007; Marder and Calabrese, 1996) because the circuit neurons and many neuromodulatory inputs are identified and characterized. The *C. borealis* STG contains 26-27 neurons (Kilman and Marder, 1996). There are about 20-25 pairs of descending modulatory neurons (Coleman et al., 1992) that contain a large number of peptide and small molecule cotransmitters (Nusbaum and Beenhakker, 2002; Nusbaum et al., 2017; Nusbaum et al., 2001) and project into the STG from the anterior oesophageal ganglion (OG) and the paired commissural ganglia (CoGs). The STG contains the essential circuitry for the generation of the relatively fast triphasic pyloric rhythm while the slower gastric mill rhythm depends on the descending inputs (Blitz and Nusbaum, 1997; Blitz and Nusbaum, 2011). When impulse traffic in the stomatogastric nerve that brings the modulatory inputs to the STG is blocked, the gastric mill rhythm stops and the pyloric rhythm markedly slows (Hamood et al., 2015; Hamood and Marder, 2015; Russell, 1979). In turn, exogenous application of a number of excitatory neuromodulators, such as the neuropeptide proctolin and muscarinic agonists such as oxotremorine, strongly activates the pyloric rhythm (Bal et al., 1994; Hooper and Marder, 1987; Marder and Eisen, 1984).

In previous work the effects of temperature were characterized on the pyloric rhythm when the descending modulatory inputs were left intact (Soofi et al., 2014; Tang et al., 2010; Tang et al., 2012). Here we first study the effects of temperature on the pyloric rhythm in the absence of neuromodulation, and then in the presence of exogenously applied neuromodulators.

STG preparations were “decentralized”, that is the action of descending modulatory inputs from anterior ganglia were removed by cutting the stomatogastric nerve (stn). Figure 1A shows the recording configuration used and the placement of the extracellular electrodes on the motor nerves used to monitor the activity of the neurons that are active in the pyloric and gastric mill rhythms. Previous work (Hamood et al., 2015) quantified the effects of the removal of the descending modulatory inputs on large populations of STGs and demonstrated a range of network outputs when STGs were decentralized. Some preparations continued to generate normal but slow pyloric rhythms, while others lost activity entirely or became arrhythmic (Fig. 1B 11 ^°^C).

**Figure 1.**
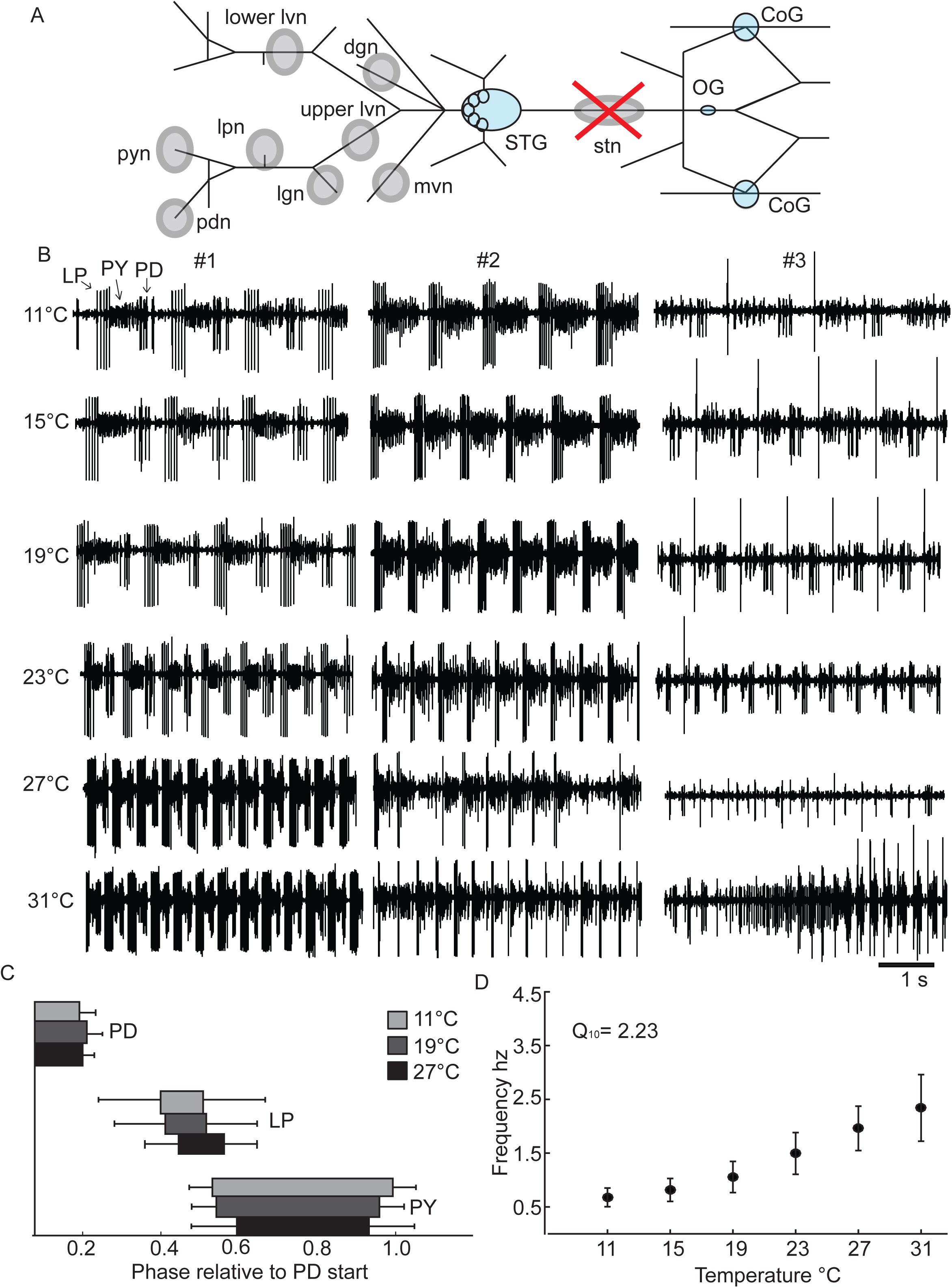
Recording configuration and control data from decentralized preparations. A. Simplified diagram of the STNS as it would appear dissected and pinned out onto a Sylgard coated petri dish. The Stomatogastric Ganglion (STG), Commissural Ganglia (CoGs) and Esophageal Ganglion (OG) are colored light blue. Grey circles indicate where Vaseline wells are made around nerves of interest for extracellular recordings. The stn (stomatogastric nerve) well with the large red ‘X” indicates where the stn is cut for decentralization after the well surrounding the stn had been filled with TTX and sucrose to silence the nerve. All the other wells, which fall posterior (left on diagram) to the STG, indicate the nerves where extracellular recordings were made. B. Five seconds of raw extracellular traces from the lvn from three different decentralized preparations (#1, #2, #3), at 11°C, 15°C, 19°C, 23°C, 27°C and 31°C. The activity shown from these preparations represents the variety of activity seen in the decentralized condition across temperature. C. The average phase relationships of the neurons that generate the triphasic rhythm for three temperatures, 11°C (light grey), 19°C (dark grey) and 27°C (black). The phase is calculated relative to the start of the PD burst per cycle. The bars represent the standard deviation of the data. Only preparations with a triphasic rhythm are included in these plots. (N@11=24, @15=24, @19=25, @23=23, @27=18, @31=11). D. The mean frequency of the rhythm calculated from the activity of the PD neurons, across temperature. The bars represent the standard deviation of the data. Only preparations with rhythmic PD bursts are included in this plot. (N@11=24, @15=24, @19=25, @23=24, @27=19, @31=14).

### The effects of temperature on the decentralized pyloric rhythm

We first characterize the effects of temperature on 40 decentralized preparations, as these data constitute baseline measurements for the effects of exogenously applied neuromodulators. Figure 1B shows examples of the triphasic pyloric rhythm as a function of temperature in preparations from three different animals. In preparation #1, we see a characteristic and classic pyloric rhythm (activity of LP, PY, and PD) at all temperatures from 11 ^°^C to 31 ^°^C. Over this temperature range the pyloric rhythm frequency increased from 0.8 Hz to 2.4 Hz. Preparation #2 also displayed a classical pyloric rhythm from 11 ^°^C to about 23 ^°^C, but when the temperature was further increased, the rhythm became less regular, and showed intermittent changes in activity. Preparation #3 shows an example of a weaker pyloric rhythm at 11 ^°^C, characterized by a single LP spike/burst. As the temperature was increased, LP activity first dropped out, and eventually the rhythm became highly irregular. These three examples show the ranges of activity routinely seen at 11 ^°^C under these conditions (Hamood et al., 2015; Hamood and Marder, 2015).

Figure 1C shows the normalized phase relationships for the activity of the LP, PY and PD neurons at different temperatures, for those stretches of data that maintained a regular pyloric rhythm. Note that these phase relationships are relatively temperature invariant, as had been reported for data from preparations with modulatory inputs intact (Tang et al., 2010). Figure 1D shows the effects of temperature on the frequency of the triphasic rhythm for those stretches of data when it was triphasic. Similar to what was seen with the modulatory inputs intact (Tang et al., 2010), the apparent Q_10_ of the frequency was 2.23.

### Characterization of different activity patterns

The raw data shown in Figure 1B illustrates that at high temperatures it is quite common to see disturbances or “crashes” in the activity patterns produced by neurons of the pyloric circuit (Tang et al., 2012). We wanted to display both the kinds of activity patterns displayed by different preparations and to characterize the potential effects of modulators on these. Because the recordings are often not stationary, but individual stretches of data show transitions between qualitatively different states of activity, we classified the raw data into eight different activity states, as defined in Figure 2. The **GLR** (lavender) (Gastric-like Rhythm) state has a robust pyloric rhythm with bursts of activity in the gastric LG neuron (pink traces) and DG neuron (not shown). The **NTf** (Normal triphasic fast) (dark blue) state is a normal triphasic rhythm with a frequency greater than 0.8 Hz. The **NTs** (Normal triphasic slow) (yellow) state is a triphasic rhythm slower than 0.8 Hz, with a concomitant change in phase (Hamood et al., 2015). The **LP01** (LP, 0 or 1 spike/burst) (orange) rhythm has normal PD, PY alternation with either 0 or 1 LP spike/burst. The **IT** (Intermittent Triphasic) (brown) rhythm is an intermittent triphasic rhythm (Fig. 2) with short periods of triphasic activity with interruptions. The **PD** (just PD bursting) (dark green) traces are associated with activity only in the PD neurons. The **AF** (Atypical Firing) (light green) state is atypical firing. The **S** (All silent) (black) state is all units silent (Fig. 2).

**Figure 2.**
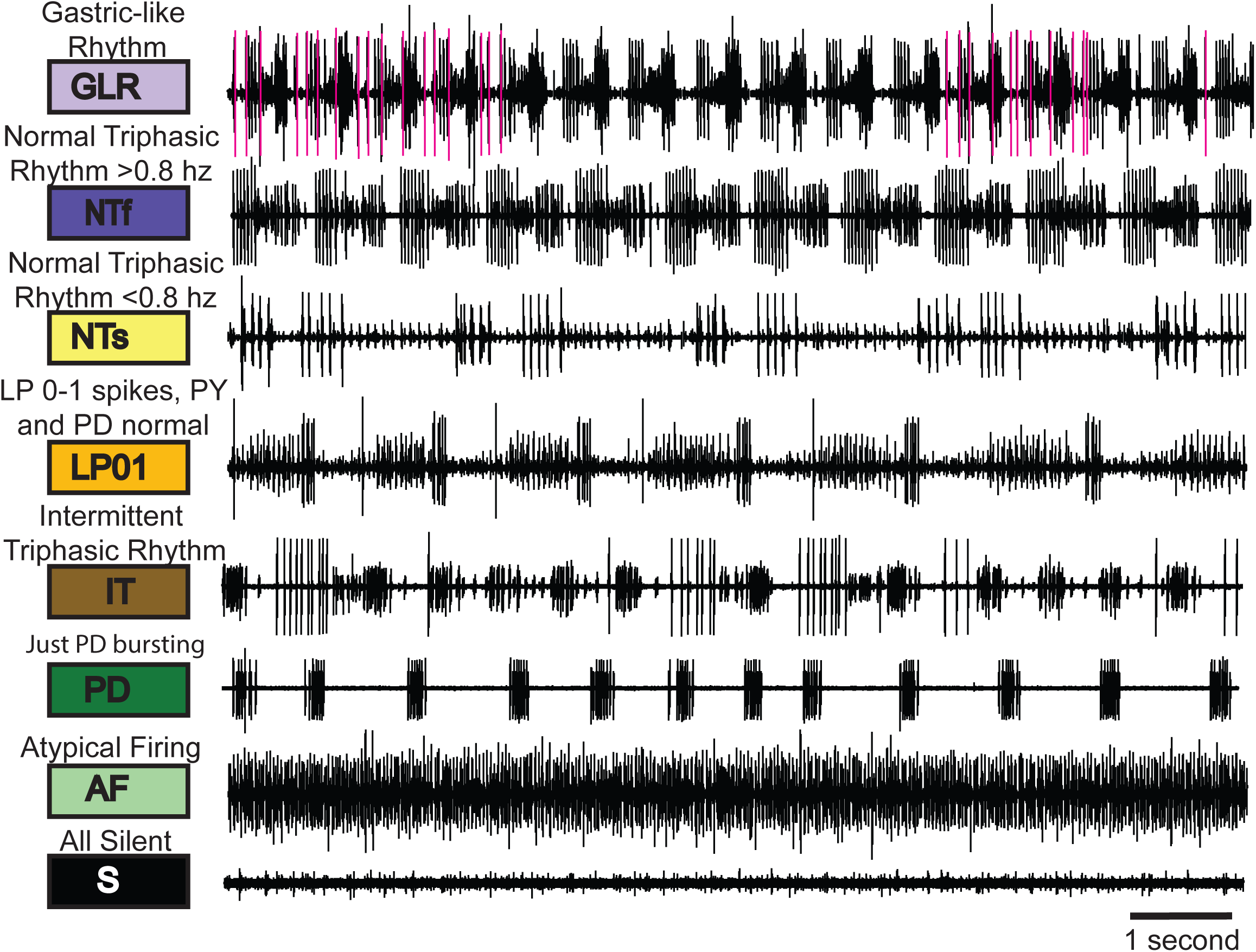
The categorization system. The eight categories are illustrated with sample raw traces that represent each category used in this analysis. The colored boxes designate the color that represents that particular category. Above each box is the descriptive name of each category and the abbreviation for each category is indicated within the colored box.

### Temperature-Dependent State Transitions

Using the classifications defined in Figure 2, we classified 150 seconds of data into state blocks, with each block at least 1 second long. This allows each stretch of data to show as many as 149 state transitions. Figure 3A shows the three preparations from Figure 1B analyzed from 11 ^°^C to 31 ^°^C in this way. Preparation #1 was slow triphasic (**NTs**) at the two lowest temperatures and **NTf** at the four higher temperatures. Preparation #2 also went from **NTs** to **NTf** at lower temperatures, but made a variety of other state transitions at higher temperatures. Preparation #3 was **LP01** at lower temperatures and again showed state transitions at higher temperatures.

**Figure 3.**
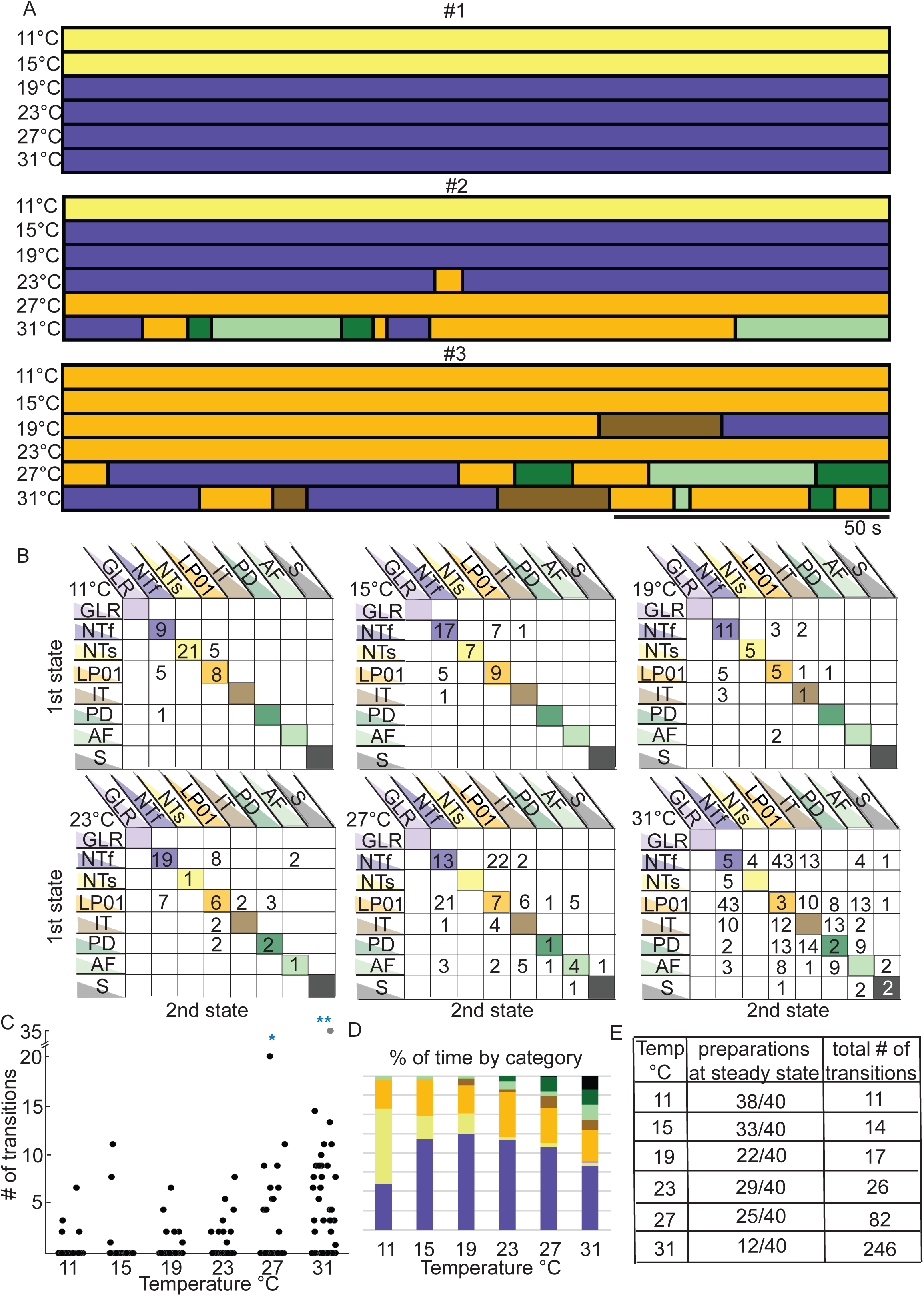
Transitions seen in decentralized preparations. A. Same preparations shown in Figure 1 (#1, #2, #3) with activity patterns of each preparation at each of the 6 temperatures, categorized as described in Figure 2. Each horizontal bar represents one data file of recordings 150s long, during which the preparation is at a stable temperature, indicated to the left. B. These matrices contain the cumulative data of transitions and stable activity patterns, for all the decentralized preparations (N=40) at each temperature. The y-axis indicates the starting activity state and the x-axis indicates the state to which the transition occurs. The diagonal lists the number of preparations that were at that particular state for an entire temperature/file, meaning they did not transition across states for that 150 s. C. Cumulative data showing the number of transitions counted per preparation (each black dot is a single preparation) as a function of temperature. The number of transitions is significantly higher at 27°C and 31°C than the other temperatures (*p<0.01 for 27°C and **p<0.001 for 31°C calculated with a one-way ANOVA adjusted for multiple comparisons). D. Each bar represents the 100% of time of all the preparations as a function of temperature. This shows that more types of activity patterns appear as the temperature is increased and more of the less stable activity patterns are present. E. The tabulated counts of preparations at one state for an entire file (steady state) and the number of transitions counted across all preparations at each temperature.

The state transitions seen at higher temperatures in Figure 3 have highly variable durations. To determine whether there are specific patterns of state transitions within a given preparation, we plotted the specific transitions shown across temperatures. There are 6 plots shown in Figure 3B, one for each temperature (indicated in the top left of each plot). The y axis shows the state of the preparation before a transition, and the x axis shows the state of the preparation after a transition. The numbers in the colored diagonals are the numbers of preparations that made no transitions. That is to say at 11 ^°^C, 9 preparations showed **NTf** throughout the entire recording time, while 21 preparations were **NTs** throughout and 8 were **LP01** throughout. The numbers in the white boxes not on the diagonals are the number of those specific transitions across all 40 preparations. That is, there were 5 LP01 transitions to **NTf**, and 5 NTs to **LP01**.

As the temperature was increased, the number of preparations on the diagonal generally decreased, and the number of transitions between other states increased. At 31 °C, there were only 12 preparations left on the diagonal and there were 246 transitions. Additionally, the destabilizing effects of temperature can be seen by the fact that the number of transitions in the lower portions of the plots increased at higher temperatures.

Figure 3C shows a plot of the total number of transitions for each preparation (each preparation is represented by a single point) at each temperature. A one-way ANOVA adjusted for multiple comparisons shows that the number of transitions at 27 °C and 31 °C were statistically different from all other temperatures. Figure 3D displays the total amount of time these preparations spent in each category, and again illustrates that temperature increases the frequency of the pyloric rhythm (the dark blue bars) but eventually at high temperatures less and less time is spent in a characteristic triphasic rhythm. Figure 3E summarizes the number of preparations that maintained steady-state activity as a function of temperature and the total number of transitions from all 40 preparations at each temperature.

### Oxotremorine and proctolin stabilize the pyloric rhythm to temperature

Muscarinic agonists and the neuropeptide proctolin have been long known to strongly activate the pyloric rhythm (Elson and Selverston, 1992; Hooper and Marder, 1987; Marder and Eisen, 1984; Marder and Paupardin-Tritsch, 1978; Nagy and Dickinson, 1983; Nusbaum and Marder, 1989a; Nusbaum and Marder, 1989b), by activating a voltage-dependent inward current, now called I_MI_ (Golowasch and Marder, 1992b; Swensen and Marder, 2000). Interestingly, although oxotremorine and proctolin have somewhat different neuronal targets in the STG, they do converge on the same final membrane current, and this is associated with increases in the amplitude of the slow wave bursts that govern the pacemaker activity of the STG networks (Golowasch and Marder, 1992b; Sharp et al., 1993; Swensen and Marder, 2000, 2001).

The raw data shown in Figure 4A,B show that both 10^-5^ M oxotremorine and 10^-6^ M proctolin stabilize the pyloric rhythm against temperature disruption. Preparations #4 and #5 are shown in oxotremorine, and preparations #6 and #7 are seen in proctolin. In these cases, the frequency of the pyloric rhythm increased as the temperature was increased, and the triphasic nature of the rhythm was maintained, even at 31 ^0^C. The stability of the rhythms is seen in the relative lack of transitions in all 4 preparations at all temperatures (Fig. 4C,D). This is independent of the fact that in modulator the starting frequencies were quite different. Figures 5A and 5B show the effect of temperature on the phase of each of the neurons in the triphasic rhythm in both oxotremorine and proctolin. All of the preparations remained triphasic in oxotremorine (n = 10) and proctolin (n = 10) at all temperatures. Figures 5C and 5D show the effect of temperature on the pyloric rhythm frequency in oxotremorine and proctolin. Note that the oxotremorine preparations increased in frequency far more than did the proctolin preparations. The apparent Q10 of the pyloric rhythm frequency in the oxotremorine condition is 2.07. The apparent Q10 of the pyloric rhythm frequency in proctolin is 1.79. Figures 5E,F show that the number of transitions counted (per individual) in the modulator conditions (red) were significantly different in comparison to the decentralized controls (black) at 27 °C and 31 °C. Figures 5G,H show that the preparations spent all of the recorded time in one of two stable conditions across the range of temperatures, whereas the same preparations, in the absence of modulators, displayed a range of activity patterns, with progressively less stable conditions present as temperature increases.

**Figure 4.**
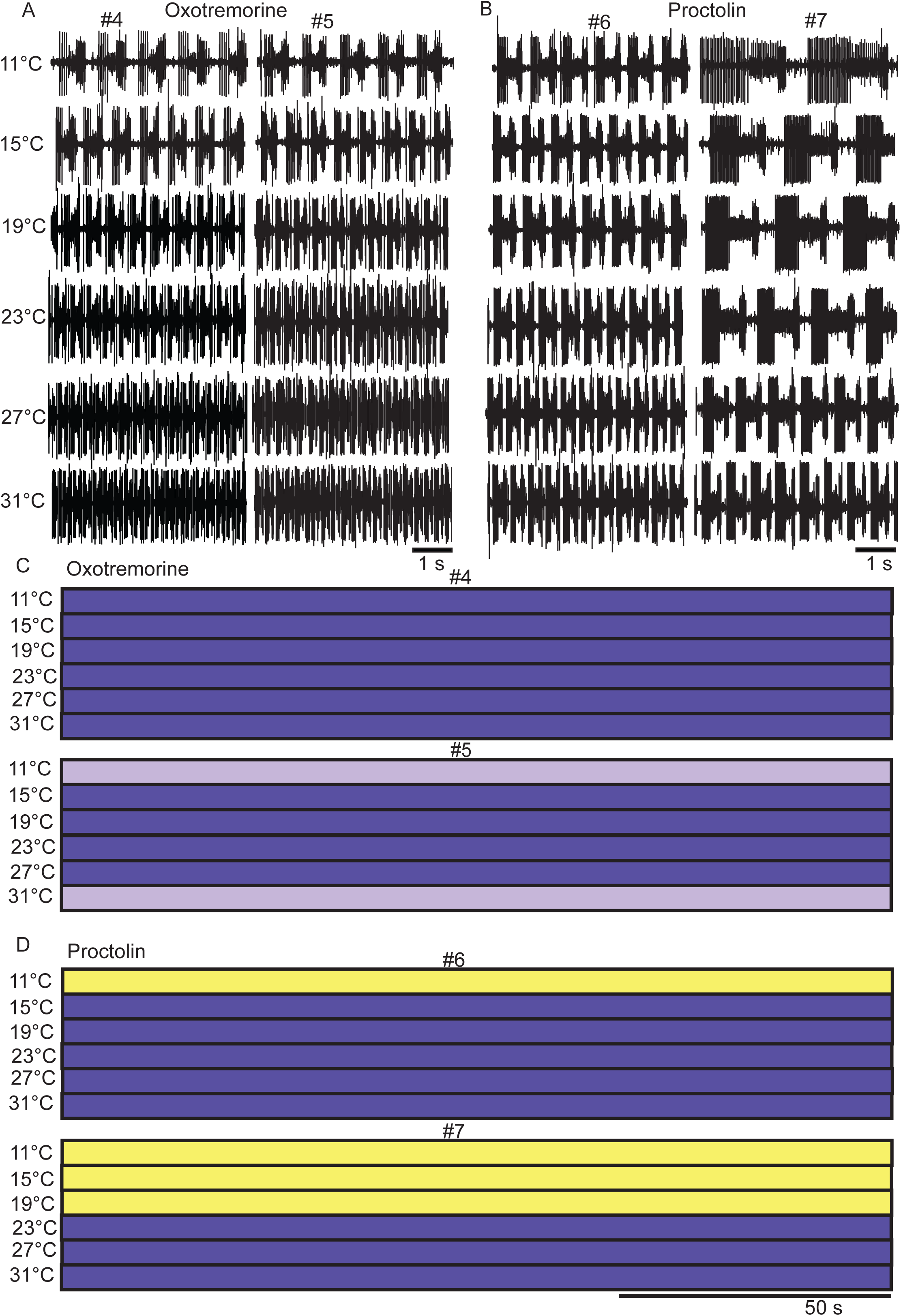
The effects of oxotremorine and proctolin as a function of temperature. A. Example raw traces from two different preparations (#4 and #5) in 10^-5^ M oxotremorine at each temperature. B. Example raw traces from two different preparations (#6 and #7) in 10^-6^ M proctolin at each temperature. C. The activity of the same preparations (#4 and #5) from panel A across the entire 150 s file as a function of temperature. D. The activity of the same preparations (#6 and #7) from panel B across the entire 150 s file as a function of temperature.

**Figure 5.**
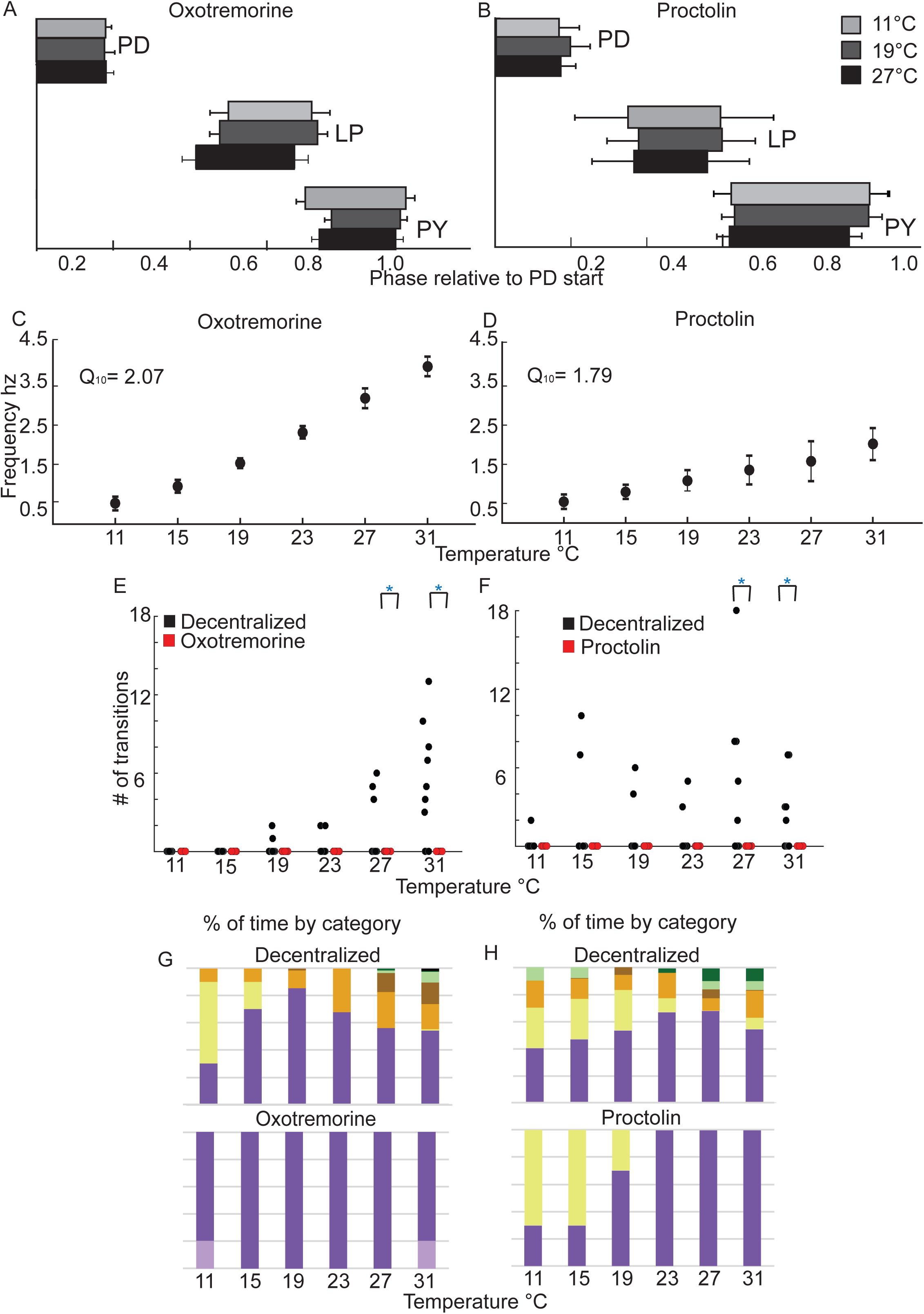
The effects of proctolin and oxotremorine on STG motor patterns as a function of temperature. A B. The average phase relationships (means with S.D.) of the neurons that generate the triphasic rhythm for three temperatures, 11°C (light grey), 19°C (dark grey) and 27°C (black) in 10^-5^ M oxotremorine (A) (N=10) and in 10^-6^ M proctolin (B) (N=10). C.D. The mean (with S.D.) frequency of the rhythm calculated from the activity of the PD neurons, across temperature in 10^-5^ M oxotremorine (C) (N=10) and in 10^-6^ M proctolin (N=10) (D). E.F. The number of transitions/preparation in the decentralized condition (black circles) and in the presence of 10^-5^ M oxotremorine and 10^-6^ M proctolin (red circles) at each temperature (N=10 for each). Decentralized and oxotremorine and proctolin conditions are significantly different at 27°C and 31°s (*p<0.01; paired t-test). G. Each bar represents the cumulative sums of the conditions for all preparations as a function of temperature where the top bar graph is the sum of the decentralized condition and the lower bar graph is the sum of activity patterns in the oxotremorine condition. This shows that more types of activity patterns appear as the temperature is increased in the decentralized condition and more of the less stable activity patterns are present. In oxotremorine all preparations maintain stable activity in one of 2 robust states. H. As (G) for proctolin.

### Serotonin destabilizes the pyloric rhythm to temperature

Serotonin is liberated from the Gastropyloric Receptor Neurons (GPR) (Katz et al., 1989; Katz and Harris-Warrick, 1989), and modulates a large number of different currents in STG neurons by activating several 5-HT receptors (Clark et al., 2004; Kiehn and Harris-Warrick, 1992a, b; Zhang and Harris-Warrick, 1994). Moreover, serotonin applications can result in highly variable changes in activity across individuals (Spitzer et al., 2008).

The across-animal variability is readily observed in the raw traces seen in Figure 6A. This figure includes raw data from 5 different preparations, to illustrate the range of activity seen in the presence of 10^-5^ M serotonin, first at 11 ^°^C, and then as a function of increased temperature. Preparations #8 and #12 were triphasic at low temperature, but lost this behavior as a function of increased temperature. Preparations #9 and #11 had gastric-like behavior at low temperatures and then lost this activity at higher temperatures. The raw traces in Figure 6A and the state-blocks in Figure 6B show that in all cases, the activity patterns seen at higher temperatures were disorganized, with a variety of different non-triphasic patterns observed.

**Figure 6.**
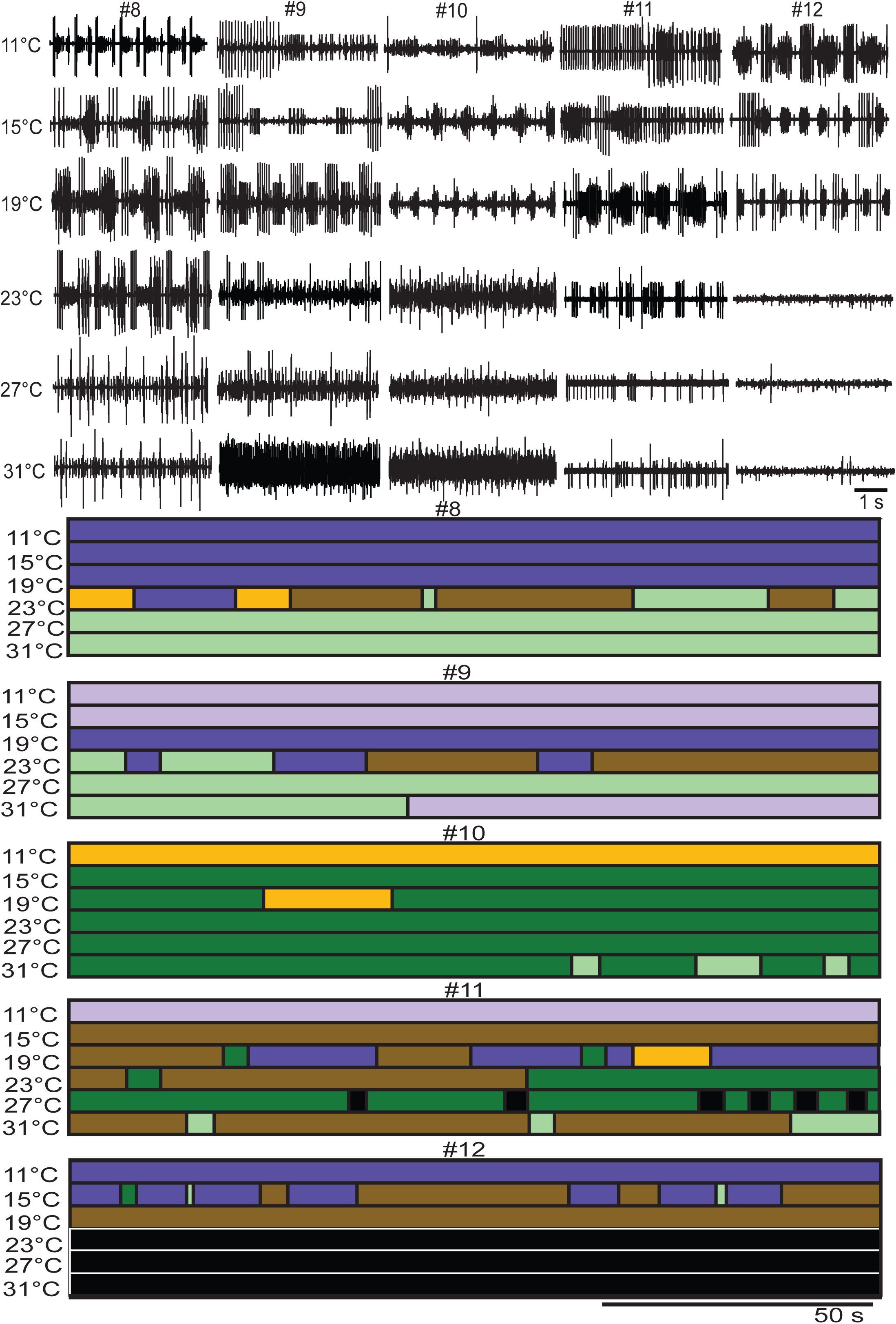
The effects of serotonin as a function of temperature. A. Example raw traces from five different preparations (#8, #9, #10, #11 and #12) in 10^-5^ M serotonin at each temperature. B. The activity of the same preparations (#8, #9, #10, #11 and #12) from panel A across the entire 150 s file as a function of temperature, using the color codes as previously described.

Figure 7A presents the state-transitions seen in serotonin for data pooled for all animals. Interestingly, all preparations are in a steady state at 11 °C (all individuals accounted for in the diagonal blocks of the diagram). The largest number of transitions appears at 19 °C when the preparations are beginning to become unstable. These preparations are significantly more temperature sensitive in the presence of serotonin than in the absence of modulation. This can be seen in Figure 7B, as the numbers of transitions per individual are significantly different by 19 °C and 23 °C. The number of transitions decreases again at higher temperatures in the presence of serotonin as preparations maintain a particular form of disrupted pyloric activity. Figure 7C shows that by 27 °C and 31°C preparations show almost no triphasic activity. Figure 7D shows the number of preparations at steady state and the total numbers of transitions present in the data in serotonin at all temperatures. The paucity of triphasic rhythms in these data sets across temperature makes it difficult to meaningfully plot and analyze either pyloric rhythm frequency or phase across temperature in serotonin.

**Figure 7.**
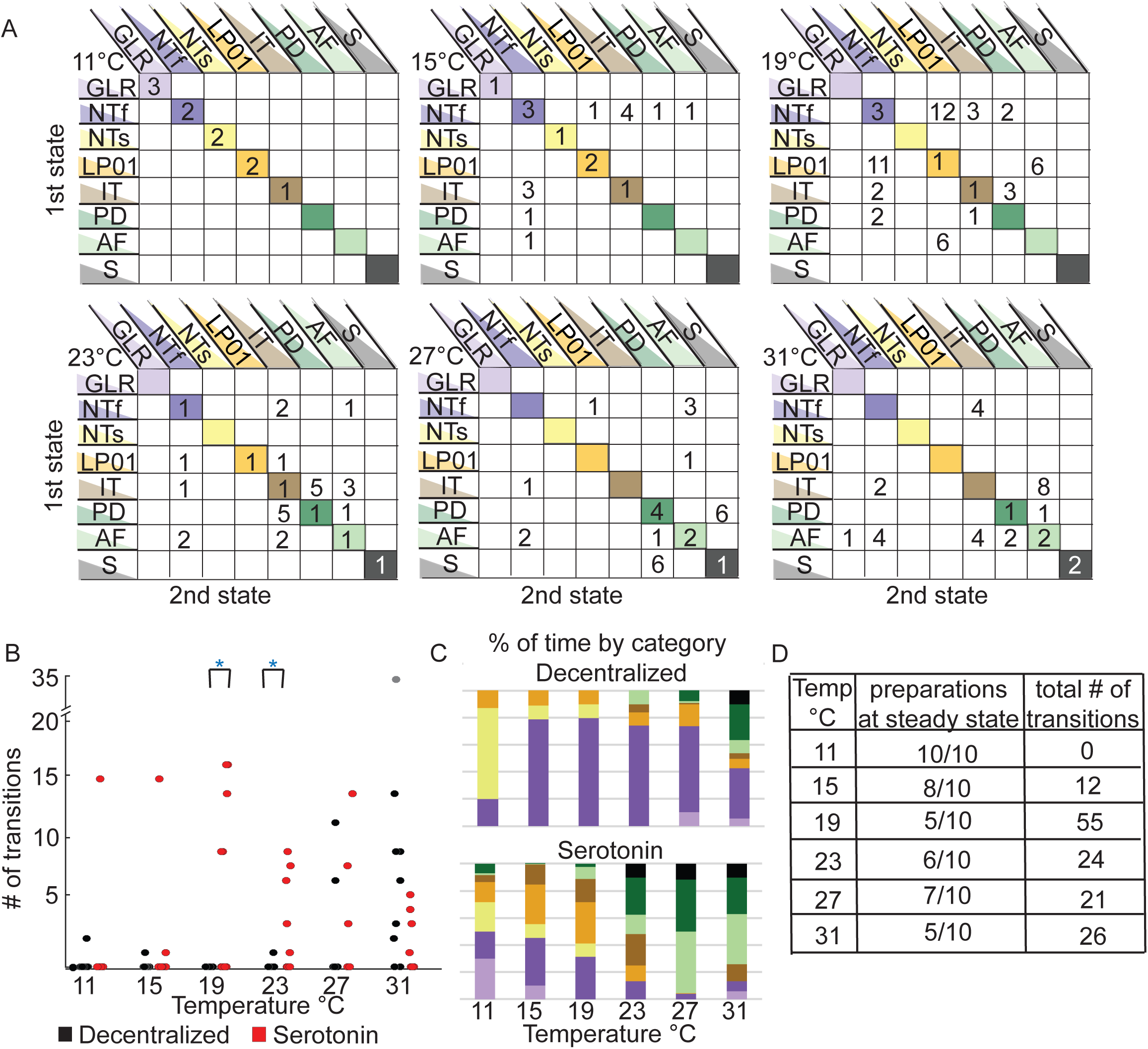
Quantification of transitions seen in serotonin across temperature. A. Matrices of the cumulative data of transitions and stable activity patterns, for all the decentralized preparations (N=10) at each temperature. The y-axis rows indicate the first activity state the preparation is in, while the x-axis columns indicate the state transitioned to. The diagonal lists the number of preparations that were at that particular state for an entire temperature/file, meaning they did not transition across states for that 150 s. B. The number of transitions per preparation in the decentralized condition (black circles) and in the presence of 10^-5^ M serotonin (red circles) at each temperature (N=10). The number of transitions per preparation in the decentralized and serotonin conditions, when each preparation is compared in a pairwise t-test the number of transitions are significantly different at 19°C and 23°s (*p<0.01). C. Each bar represents the cumulative sums of the conditions that all the preparations were in as a function of temperature where the top bar graph is the sum of the decentralized condition and the lower bar graph is the sum of activity patterns in present in the serotonin condition. This shows that more types of activity patterns appear at lower temperatures in serotonin and as temperature is increased, more of the less stable activity patterns are present in both conditions. D. The tabulated counts of preparations at one state for an entire file (steady state) and the number of transitions counted across all preparations at each temperature in the serotonin condition.

## DISCUSSION

A fundamental question in neuroscience is understanding how reliable and robust circuit outputs can result from variable components. This has been illuminated by the finding in both theoretical and experimental work that there can be multiple and degenerate solutions to the production of similar outputs (Britton et al., 2013; Goaillard et al., 2009; Gutierrez et al., 2013; Muszkiewicz et al., 2016; Prinz et al., 2004). Therefore, in principle, one would expect to find that individuals of the same specie could show considerable variability in the parameters that govern intrinsic excitability, synaptic strength, and the details of neuronal structure across animals. In fact, across animals there appears to be a 2-6 fold range in synaptic strengths, conductance densities, and ion channel mRNA expression (Golowasch et al., 1999; Roffman et al., 2012; Schulz et al., 2006; Schulz et al., 2007). Moreover, the same identified neuron or neurons in different animals can show variable branching structures (Bucher et al., 2007; Hay et al., 2013; Otopalik et al., 2017).

Even if individual animals show similar motor patterns, to the extent that they do so with different sets of membrane and synaptic conductances, one would expect that extreme perturbations will reveal the consequences of those differences. In previous work we used temperature as a perturbation to probe the animal-to animal differences in conductance densities (Rinberg et al., 2013; Tang et al., 2010; Tang et al., 2012), and showed that the pyloric rhythm and its isolated pacemaker neurons from individual animals “crashed” with individual dynamics. These experiments were done with the descending modulatory inputs from the CoGs and OG left intact, which means that the recorded pyloric rhythms were enhanced by the release of endogenous neuromodulatory substances. In the absence of these substances, the pyloric rhythm is less robust, slows or stops, and becomes less regular (Hamood et al., 2015; Hamood and Marder, 2015), and one might expect that these preparations would be more sensitive to extreme temperatures. In this paper we confirm that decentralized preparations, in which the modulatory inputs are removed, are variable at 11 ^°^C, and become even more so at higher temperatures. Thus, in the absence of modulatory inputs, temperature perturbation effectively reveals the consequence of underlying differences in network parameters across animals.

### Classification of Network States

Tang et al (2012) attempted to design a measure of the extent to which a given set of recordings become disorganized or crashed, and developed a robustness index. The robustness index was only partially successful in capturing the essence of the fluctuating and intermittent patterns of activity that are seen in the raw data (e.g. Fig. 1). We experimented with a variety of other quantitative data analysis methods, but ran into several problems. A) Once the preparation reached a temperature at which it starts to become disrupted, it goes through patterns of state transitions of widely different durations and characteristics. This means that collapsing the data via spectral analyses or other common ways of analyzing rhythmic data, fail to capture some of the qualitatively interesting and significant features of the data. B) These are long experiments with many conditions (multiple temperature ramps in the presence and absence of modulator). Therefore, we were restricted in the amount of time we were able to record each condition, and settled on analyses of 150 seconds after the preparation had reached a given temperature. Given the period of the pyloric rhythm, and the durations and numbers of qualitatively different states in the raw data, this is not enough data with which to use machine learning or other automated methods of data classification. This problem is exacerbated by the lack of obvious stationarity shown at any given temperature.

After unsatisfactory attempts to analyze these data in a way that preserves its qualitatively rich properties, we settled on using the best pattern classifier we know for these sorts of data: the human observer. Subsequently, we have encountered numerous other investigators with similar problems in which an experienced human observer can classify data using multiple features without a struggle, even in relatively small data sets. In this paper we defined 8 qualitatively different network states after looking at large amounts of raw data. This is a balance between using very broad classifications that might lead to fewer states or narrower classifications that might lead to the identification of many more states. Again, these decisions were made on the basis of experience with the stomatogastric motor patterns and experience-dependent evaluation of what is likely to be important for this network. We would argue that accrued knowledge by investigators is not necessarily a bad way to design a classification scheme, as long as this is done transparently, and not hidden in decisions about algorithm choices.

Unfortunately, because each stretch of data was relatively short, we are unable to determine whether there are specific sets of preferred transitions among states, and whether the statistics of preferred transitions might differ across animals and across modulatory condition. To address this question in the future would require collecting much longer sets of data while holding the preparation at an appropriate high temperature, without damaging it.

### Modulators as Stabilizers or Destabilizers

Decentralization, which removes the effects of most of the neuromodulatory substances that routinely reach the STG, destabilizes the pyloric rhythm and produces state-transitions between a variety of outputs. This argues that the modulatory inputs to the STG may be protecting the pyloric rhythm from the deleterious effects of high temperatures. Consistent with this idea are previous studies on lobster neuromuscular junctions that showed that dopamine (Thuma et al., 2013) and serotonin (Hamilton et al., 2007) extend the temperature range at which of functional muscle and neuromuscular junction performance. In a fascinating study, Zhurov and Brezina (2005) showed that while increased temperature decreased transmitter release at an Aplysia neuromuscular junction, the modulators themselves enhanced muscle contractility in a way that compensated for this decrease, and maintained muscle activity at high temperatures (Zhurov and Brezina, 2005). A similar conclusion was suggested by a modest increase by modulators in the temperatures over which the gastric mill rhythm is active (Stadele et al., 2015). Moreover, work in *C. elegans* argues that neuromodulation is important in allowing animals to maintain appropriate thermosensory behaviors (Beverly et al., 2011). Therefore, we were curious to see whether specific neuromodulatory substances would alter the effective operating range of the pyloric rhythm in response to extreme temperature perturbation.

### The stabilizing action of modulators that activate I_MI_

Proctolin and oxotemorine activate a current called I_MI_ which is a voltage-dependent inward current that is blocked by extracellular Ca^2+^ (Golowasch and Marder, 1992b; Gray et al., 2017; Gray and Golowasch, 2016; Swensen and Marder, 2000, 2001). I_MI_ ‘s voltage-dependence makes it ideally suited to enhance oscillations (Sharp et al., 1993) because it provides an additional inward current at more depolarized membrane potentials and is blocked at the trough of the oscillation. Consequently, agonists that activate I_MI_ characteristically strongly enhance oscillatory properties of pyloric network neurons (Hooper and Marder, 1987; Sharp et al., 1993; Swensen and Marder, 2000). Thus, our expectation was that both oxotremorine and proctolin might counteract some of the decrease in membrane resistance that commonly occurs with increased temperature (Stadele et al., 2015; Tang et al., 2010), and stretch the operating range of the pyloric rhythm by enhancing the neurons’ ability to burst.

Other cell intrinsic and network characteristics can also contribute to network stabilization by these modulators. The pyloric circuit contains multiple instances of reciprocal inhibitory connections, and rebound from inhibition is an important determinant of when the follower neurons will fire (Goaillard et al., 2010; Harris-Warrick et al., 1995; Hartline, 1979). The rebound from inhibition depends both on the strength and time course of the inhibition and the rebound membrane currents that contribute, to the rebound (Sharp et al., 1996). A number of modulators that target I_MI_, including proctolin (Oh et al., 2012; Zhao et al., 2011) and RPCH (Thirumalai et al., 2006) enhance synaptic strength in pyloric neurons, and therefore will enhance the reciprocal inhibitory connections that promote rhythmic activity in the pyloric rhythm.

Additionally, the increase in the number of spikes/burst and enhancement of the amplitude of slow wave oscillation of a presynaptic neuron, characteristic of I_MI_ activation (Hooper and Marder, 1987; Sharp et al., 1993), can further increase the inhibitory input onto its postsynaptic targets, because the chemical synapses within the STG function through graded synaptic transmission (Graubard, 1978; Graubard et al., 1980; Manor et al., 1997). The neurons that generate the pyloric rhythm have hyperpolarization-activated rebound currents (I_h_) (Golowasch and Marder, 1992a; Kiehn and Harris-Warrick, 1992a). An increase in inhibitory input increases the activation of I_h_ and the rebound burst in the follower neuron. Therefore, the activation of I_MI_ on any pyloric neuron can enhance the robustness of bursting activity of multiple neurons within the network.

It is possible that the increased robustness of rhythmic circuit output, observed by the activation of a current like I_MI_, is a common solution found in many organisms. The I-V curve of I_MI_ is similar to that of the NMDA-activated current which can activate and regularize the rhythmic oscillations of motor neurons in the lamprey spinal cord (Sigvardt et al., 1985). This suggests that the NMDA receptor may contribute to stabilizing rhythmic oscillations when lampreys or other cold-blooded vertebrates encounter temperature changes.

### Serotonin as a Circuit Destabilizer

Serotonin acts on numerous receptors to modulate a variety of voltage-dependent currents in STG and other crustacean neurons (Clark et al., 2004; Kiehn and Harris-Warrick, 1992a, b; Sosa et al., 2004; Spitzer et al., 2008; Zhang and Harris-Warrick, 1994). The variability of serotonin’s actions on the decentralized *C. borealis* rhythm is shown in the 11 ^°^C data on Figure 6. Notice, that even at low temperatures the preparation to preparation variability of the decentralized preparations is higher in serotonin than in controls. We hypothesize that this occurs because serotonin modulates so many different currents, that it is likely that the starting set of conductances in each individual will lead to considerable variation in serotonin’s actions at 11 ^°^C. This individual variability seen in serotonin is then amplified when the temperature is changed.

Under the conditions used here, serotonin was a destabilizing factor, and therefore was a useful probe to reveal cryptic animal-to-animal differences in underlying intrinsic and synaptic parameters. But in the animal, serotonin is both a circulating hormone (Kravitz et al., 1980), and therefore presumably acting at low concentrations, as well as a cotransmitter released from the Gastro-Pyloric (GPR) sensory neurons (Katz et al., 1989; Katz and Harris-Warrick, 1989). A considerable body of work on dopamine in the lobster, *Panulirus interruptus* shows that low concentrations of tonic dopamine can alter the effects of higher phasic dopamine applications (Krenz et al., 2013; Krenz et al., 2014; Rodgers et al., 2013). In a similar way, it is possible that low hormonal concentrations of serotonin *in vivo*, acting in a richer modulatory environment, might have different effects on circuit stability than seen here. Moreover, serotonin enhances the operating range of the heart neuromuscular junctions in response to temperature fluctuations (Hamilton et al., 2007) in *Homarus americanus*, as does dopamine for stomach muscles in *Panulirus interruptus* (Thuma et al., 2013).

One of the puzzling features of the stomatogastric ganglion is that it is modulated by more than 30 different substances (Marder, 2012; Marder and Bucher, 2007; Marder et al., 2014). Early work on STG modulation focused on the potential of these modulators to promote behavioral flexibility (Harris-Warrick and Marder, 1991; Hooper and Marder, 1984; Marder and Bucher, 2007). Nonetheless, it is likely that much of the modulation of the STG may be physiologically necessary to maintain stable function to counteract environmental perturbations such as temperature changes. As crabs and lobsters must deal with a variety of environmental perturbations such as alterations in O_2_ tension (Clemens et al., 1999; Clemens et al., 1998; Clemens et al., 2001), pH, salinity, etc. in addition to temperature, it is possible that different subsets of modulators are called into play in response to each of these potential stressors. As some of these environmental changes are potentially linked as pH and O_2_ tension change as a function of temperature, it will be interesting to determine how multiple modulators may coordinately regulate the central pattern generating circuits of the stomatogastric nervous system to maintain the animal’s feeding behavior over a large range of environmental circumstances.

## Author Contributions

SAH and EM designed the experiments and wrote the manuscript. SAH performed the experiments, data analysis and made the figures.

## Acknowledgments

Research Supported by R01 NS017813, R01 NS081013 and R35 NS097343. We thank Drs. Michael P. Nusbaum and Farzan Nadim for comments on the manuscript.

## Methods

### Animals

Adult male Jonah Crabs, *Cancer borealis*, were purchased from Commercial Lobster (Boston, MA). Animals were housed in tanks containing artificial saline maintained at 11°C until used. Animals used in these experiments obtained between February 2013 and January 2017.

### Solutions

*C. borealis* physiological saline was composed of (in mM): NaCl, 440; KCl, 13; CaCl_2_, 26; MgCl_2,_ 11; Trizma base, 11; maleic acid, 5, pH 7.4-7.5 at room temperature. Microelectrode solution was composed of (in mM): Na_2_SO_4_, 15; NaCl, 20; MgCl_2_, 10; K gluconate, 400 (Hooper et al., 2015). All chemicals were obtained from Sigma Aldrich.

### Modulators

Proctolin 10^-6^ M (Bachem), oxotremorine 10^-5^ M (Sigma Aldrich) and serotonin 10^-5^ M (Sigma Aldrich) were dissolved in physiological saline and applied through a continuously flowing superfusion system running at approximately 500 ml/hour.

### Electrophysiology

Stomatogastric nervous systems were dissected from crab stomachs and pinned out onto Sylgard coated petri dishes. Electrically isolating Vaseline wells were constructed around nerves of interest for extracellular recordings. For most preparations, these nerves were the upper LVN, the lower LVN, PDN, PYN, LPN, DGN, LGN and MVN. Another well was built around the STN to later be filled with TTX to block impulse traffic in the STN. The recordings were made with stainless steel pin electrodes and A-M systems amplifiers. The amplifiers were connected to a DigiData 1440A which was further connected directly to a computer. The data were recorded using pClamp software (Axon instruments). Each file collected 150s of data. The superfused physiological saline was passed through a peltier device (Warner Instruments) which maintained temperature at 11°, 15°, 19°, 23°, 27° or 31°C during temperature ramps. While recording was continuous throughout the entire duration of the experiment, the data analyzed in this paper were taken when the temperature was stable (within +/-0.3 degrees Celsius) at one of the previous listed temperatures. Each preparation was subjected to three temperature ramps. The duration of each temperature ramp was ∼45 minutes long. The first was with the descending modulatory inputs intact. In the second ramp, the descending nerves were first silenced by TTX 10^-7^ M (Sigma Aldrich) being added to a well surrounding the stomatogastric nerve which was then cut. The preparation was left at 11°C for 30-45 minutes before the subsequent temperature ramp. In the third ramp, one of the previous listed modulators (oxotremorine, proctolin or serotonin) was added to the superfused saline. The modulator temperature ramp was started after ∼5 minutes (for oxotremorine and proctolin) or ∼15-25 minutes (for serotonin) after the saline containing the modulator was circulating through the peltier, to make sure the bath saline was completely replaced with the modulator saline and physiological effects of the modulator were seen.

### Analysis

Data were analyzed using Spike2 v 7.0 (Cambridge Electronic Design) and MATLAB v R2012b (Mathworks). MATLAB and SigmaPlot v 11.0 (Jandel Scientific) were used for statistical analyses. After visual inspection of long stretches of data from many conditions, we described the states of activity as 1 of 8 categories. Each data set was then manually scored, across time, temperature and modulatory condition, with the behavior of one of these categories. Transitions from one category to another shorter than one second, were not noted. Subsequently, transition data sets were counted for number of transitions, pair-wise transitions present and their frequency and how long preparations exhibited the denoted behavior.

### Statistics

Paired T-tests were used to calculate significance (p<0.01*, p<0.001**) on the number of transitions counted for each preparation, at each temperature in the decentralized vs. modulator conditions. Kruskal-Wallis one-way ANOVA, adjusted for multiple comparisons, was used to calculate significance in the number of transitions counted for each preparation across the 6 temperatures in the decentralized condition. Matlab was used to perform these calculations.

